# More the merrier? Influence of Mango Orchards on the Composition of the Reptile Communities of the Lateritic Plateaus, Maharashtra, India

**DOI:** 10.1101/2024.04.21.590433

**Authors:** Prathamesh Amberkar, Rahul Mungikar

## Abstract

Change in land use patterns affects a landscape by changing the composition of local flora and fauna. The lateritic plateaus in Konkan are one such example of this rapid land use change. The area between the West coast and the Northern-Western Ghats of India, the Konkan region is characterized by these plateaus. Although these plateaus are rich in endemic flora and fauna, they are considered as wastelands due to lack of woody vegetation and thus, are subjected to various land uses. Mango cultivation in this region has become a major source of income since it’s known for its Alphonso variety of mangoes. Hence, over the last few decades, many plateaus have been converted into mango orchards. However, how the conversion of plateaus to mango orchards have effected species composition of different groups of organisms remains unclear. In the present study, we sampled the lateritic plateaus and plateaus converted into mango orchards in Ratnagiri, Maharashtra, to investigate the response of the reptile communities. We observed more species heterogeneity in the orchards compared to the plateaus. Since, the orchards offered more diverse microhabitats for species to occupy. Moreover, it appeared that there was a species turnover when the plateaus were converted into orchards. The generalist and widespread species such as agamids and skinks and very few specialist species were comparatively more abundant in the orchards. On a contrary, the plateaus without orchards harboured more specialist species like *Hemidactylus albofasciatus*, *Echis carinatus* and *Ophisops jerdonii* compared to the generalist species.

## 1. INTRODUCTION

The escalating human population over the course of several millennia has played a pivotal role in reshaping the Earth’s natural environment (See also. Lambin et al., 2001). The transformation of pristine habitats into agricultural lands and urban spaces has emerged as a primary driver behind the alarming decline in global biodiversity and the extinction of various species. This alteration when aggregated, significantly affects the key aspects of ecosystems. While extensive global studies have investigated the impact of land use change on biodiversity (Ojima et al., 1994), it is imperative to delve into this issue at a local scale for more effective conservation management. Although tropical regions harbour species richness, there has been a strong bias toward geographic locations, with more studies on impact of land use change on biodiversity being carried out in Europe and North America, compared to the tropics. Moreover, there exists a bias towards studying the impacts of land use changes on forest ecosystems over open ecosystems like grasslands and savannas, which are also integral to biodiversity (Davison et al., 2021).

Open ecosystems are one of the most threatened habitats supporting unique biodiversity. These ecosystems are either considered degraded forests or wastelands and are vulnerable to land use changes. The western coastal region of Maharashtra, India, known as Konkan, is characterized by such open ecosystems, the lateritic plateaus, locally known as ‘*Sadas*’. Since these plateaus have been eroded over millions of years (Widdowson and Cox, 1996), there’s very little soil present here. Majority of the land is exposed and devoid of woody plant species. The herbaceous vegetation on plateaus goes through seasonal successions and appears to be barren during summers and winters (Lekhak and Yadav, 2012), and hence considered as wastelands. The ‘wasteland’ status has been exploited and the lateritic plateaus have been subjected to various land uses (Watve, 2013). The district of Ratnagiri in Maharashtra, India is famous for the Alphonso variety of mangoes and is the leading producer in Maharashtra (Government of India et al., 2018). To achieve this level of mango production, the locals, over the past few decades, have started converting these plateaus into mango orchards (Burondkar et al., 2018).

Conversion of natural habitat into agriculture might have varying effects on biodiversity depending on the type of agriculture, geographic location and functionality of the taxa (Newbold et al., 2020). Among vertebrates, birds and mammals have received more attention, leaving a significant gap in our understanding of the impact of habitat conversion on reptiles (Davison et al., 2021). Reptiles, despite having more described species than mammals and amphibians, have been remarkably understudied concerning land use change.

The reptile species inhabiting the plateaus, like the White-striped Viper gecko (*Hemidactylus albofasciatus* Grandison & Soman, 1963)(Mirza and Sanap, 2012), Jerdon’s snake-eyed lizard (*Ophisops jerdonii* Blyth, 1853), Beddome’s snake-eyed lizard (*O. beddomeii* Jerdon, 1870) and Saw-scaled viper (*Echis carinatus* Schneider, 1801) (Gaikwad et al., 2009) are more prevalent in open and dry habitats like these plateaus, with *H. albofasciatus* endemic to these plateaus (IUCN, 2011). The response of species like *H. albofasciatus* and *E. carinatus*, at a population level, to the conversion of plateaus to orchards is already known. These species are more prevalent in undisturbed sites compared to orchards (Jithin et al., 2023). The response of reptiles to land use change at a species level might be different compared to the community level. At a community level, there can be a species turnover, with some agriculture types filtering out specialist species and harbouring more generalist species (VegalAgavo et al., 2021) or vice-versa. Making it crucial to assess community-level response as well. Hence, we examined the influence of the conversion of lateritic plateaus to mango orchards on the reptile communities of the lateritic plateaus.

Reptiles are ectotherms, i.e. they are affected by the thermal and biophysical environment that they experience. Ectotherms physiology and ecology are intimately linked. Different microhabitats provide different biophysical conditions. Hence, microhabitats play a crucial role in the survival of reptile species, and their availability is a significant factor in determining reptile presence in a landscape (Scheffers et al., 2014). Reptiles on the lateritic plateaus utilize the empty spaces under rocks as refuges during the day, highlighting the importance of these microhabitats for their survival (Gaikwad et al., 2009; Mirza and Sanap, 2012). Converting the plateaus into orchards might alter the existing microhabitats (Jithin et al., 2023). Therefore, an examination of the microhabitats occupied by reptiles in both plateaus and orchards is essential. Hence, we looked at the microhabitats occupied by reptiles in both, plateaus and orchards.

On these plateaus, the hard rock surface does not promote the growth of any woody species of trees. Hence, the hard rock mango planting technique is used to cultivate mangoes in Konkan. The hard rocky surface is blasted with the help of explosives and the pits formed are filled by non-native soft soil from adjacent areas. Mounds are made around individual plants from the rocks excavated after blasting (Bhattacharyya et al., 2019). Hence, substratum sampling was also carried out to see if this plantation technique changes the substratum.

In conclusion, overall studies revealed that how land use change to mango cultivation in the plateaus is influencing reptile communities in the lateritic plateaus. During the entire study reptiles were sampled and microhabitat features quantified across two habitat types, the lateritic plateaus and the mango orchards, to test whether reptile diversity and composition change significantly between the lateritic plateaus and the mango orchards.

## 2. METHODOLOGY

### 2.1. Study Area

The study was conducted in Rajapur administrative block in Ratnagiri district of Maharashtra, India. Rajapur lies in the Konkan region between in the Western-Ghats and shares boundaries with the Arabian Sea towards the west. The lateritic plateaus here are at an elevation of 50-200 m (MSL) and are fragmented by rivers and valleys flowing between them. These lateritic plateaus are a mosaic of built-up areas, mango and cashew plantations, agricultural land and stone quarries. Grasses are the dominant vegetation, with no woody plant species. The herbaceous vegetation present here undergo seasonal succession and are abundant during monsoon, growing in small ephemeral pools (Lekhak and Yadav, 2012; Watve, 2013). The climate during the monsoon is warm, windy and overcast and the dry months are hot with clear sky. The temperature here varies from 22^0^C – 32^0^ C.

A total of six sites (Fig. 1) were selected for the study, three on the plateaus which were not subjected to any land-use changes (referred as ‘plateaus’ hereafter) and three sites which were plateaus earlier but are now converted into mango orchards (referred as ‘orchards’ hereafter). All the sites were at a minimum distance of 5 km from each other and were around 0.3 sq. km in area. The outline of each of these study sites was demarcated using the ‘polygon feature’ using Google Earth Pro (Supplementary figure 1).

**Figure 1:**
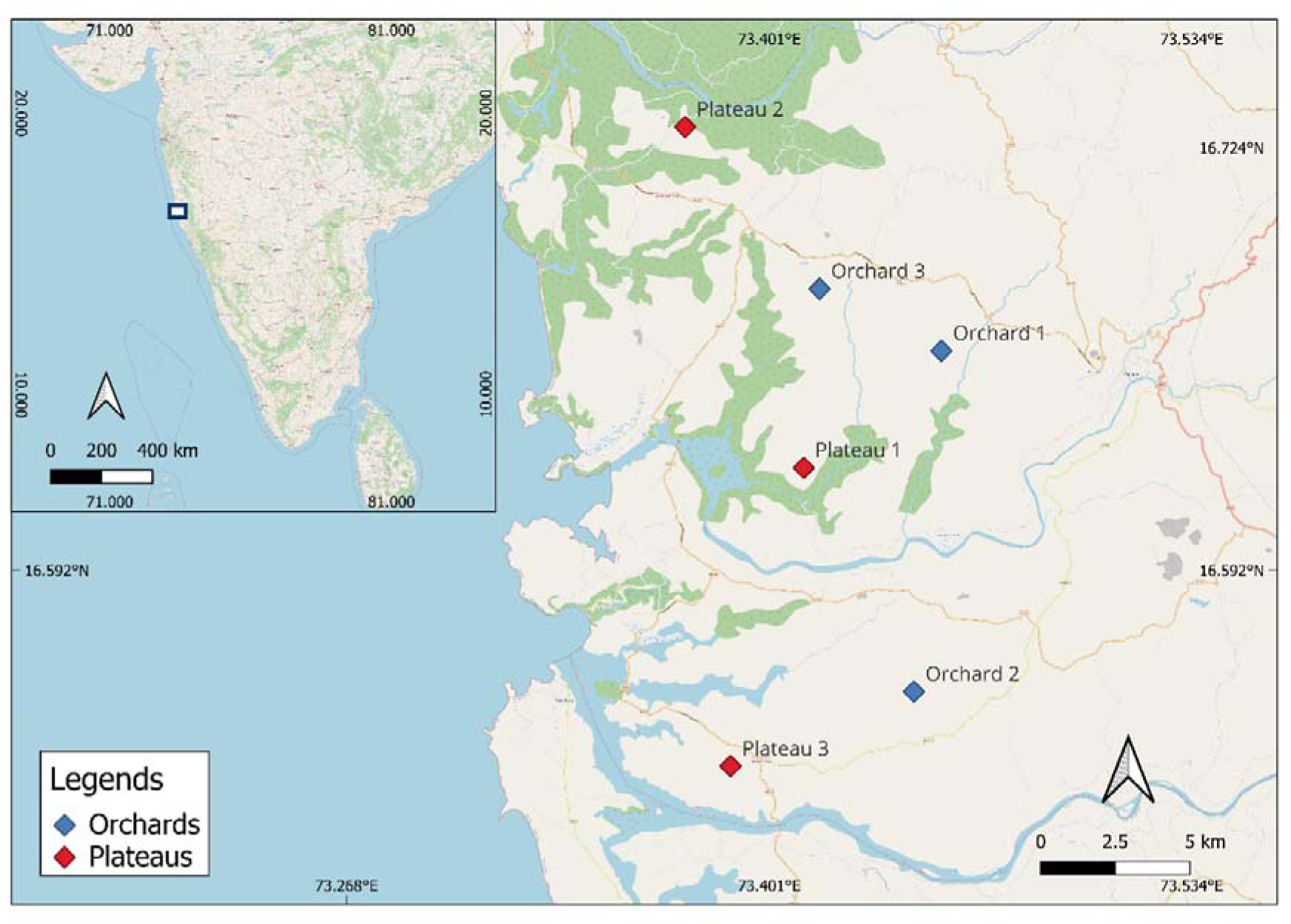
Study area map.

### 2.2. Species composition and microhabitat association

Within each site, six - eight belt transects of 100m x 6m were marked, each transect being about 100 m– 200 m apart from each other. Each transect was denoted virtually as a line feature using Google Earth Pro (Supplementary figure 1). The sampling was carried out in between September to November 2021 from 1500 hrs – 1800 hrs. Active search for reptiles was carried out along the transects and reptiles were actively looked for in the microhabitats (Dodd, 2016). The reptiles species spotted within the transect were recorded along with the microhabitats in which they were found at the time of spotting. The microhabitats were pre-classified based on a pilot study into seven main categories as follows: i) under a rock in bare patch ii) under in rock in grass patch iii) bare/open patch iv) leaf litter v) rock crevices vi) on the rocks vii) bushes/tree branches. All species were identified up to species level using the available identification keys. Species that couldn’t be identified on the filed were photographed were later identified up to the smallest taxonomic unit possible.

### 2.3. Percentage of substratum cover

To test whether conversion of lateritic plateaus into orchards changes the available ground microhabitat, further categorization of the substratum was done on the basis of the ground covering component like leaf litter, vegetation etc. The sub-categories defined as a) Exposed habitat b) Covered with leaf litter c) Covered with seasonal grasses and d) Covered with herbs and shrubs. The substrate type for each site was calculated as percentage cover within 1×1 m plots, which were randomly sampled. At each of the six site, we placed 10 plots (n = 60). Each plot was photographed using a Nikon D7500 camera. These images were later used to calculate the percentage cover of each substratum category in each plot using image analysis to test whether there was a significant difference between the substrates available for reptiles (Supplementary image 2).

### 2.4. Data analysis

All the analyses were carried out in R studio (version 4.3.1). The heterogeneity of species composition in both habitats was calculated using Shannon’s entropy (*H’*) using the ‘*vegan*’ package. To check if the *H’* values were significantly different, we performed Hutcheson’s t-test (Hutcheson, 1970), using the ‘*Hutcheson_t_test’* function in the ‘*ecolTest’* package (Salinas and Ramirez-Delgado, 2021). Additionally, we calculated *H’* to assess the heterogeneity of microhabitats occupied by reptiles in each habitat type. Multinomial species classification method (Chazdon et al., 2011) was used to classify species into habitat specialist and generalist using the ‘*clamtest*’ function in ‘*vegan*’ package. The package classified the species into three categories 1) Specialist plateaus (referred as ‘specialists’ hereafter) 2) Specialist orchards (referred as ‘generalist’ hereafter) and 3) Too rare (species with less observations).

Whittaker’s beta-diversity (Whittaker, 1972) was calculated across sites using the *’betadiver’* function in the ‘*vegan’* package to examine the differences in species composition amongst sites. The dissimilarity matrix was represented using hierarchical clustering, using the ‘*hclust*’ function. We used Chi-squared test to evaluate whether there was a significant dependency between the abundance of species across sites using the ‘*chisq.test*’ function in R using the ‘*vegan*’ package. Later, to visualize the distribution of species in each site, we used Correspondence Analysis (CA) through the ‘*CA*’ function in the ‘*FactoMinerR’* package in R (Husson et al., 2023). Similarly, we used the CA to visualize the relationship between microhabitats and reptile species.

To estimate percentage of substrate cover across habitats, images of each 1×1 m plot were analysed using the *ImageJ* software (Schneider et al., 2012). The colour threshold of each substrate category was adjusted manually and then the image was converted into a binary image (Supplementary figure 2). The percentage cover of each of the plots was calculated using the ‘measure’ tool in *ImageJ* based on the pre-classified categories (Broeke et al., 2015). Kruskal-Wallis test using the ‘*rstatix*’ package and followed by the Dunne’s post-hoc test (Kassambara, 2023) was used to test difference in the substratum across sites. The plots were visualised using the ‘*ggbetweenstats*’ function in the ‘*ggstatsplot*’ package (Patil, 2021).

## 3. RESULTS

### 3.1. Composition of reptile species

We sampled a total of six sites (three mango orchards and three plateaus) and found 10 species of reptiles in both habitats (Table 1). Species like *H. albofasciatus* (n = 61), *H. cf. varadgirii* (n = 7), *E. carinatus* (n = 15) and *O. jerdonii* (n = 45) were more prevalent in the plateaus compared to the orchards. The Orchards were more abundant with skinks like *E. carinata* (n = 16) and *E. allapallaensis* (n = 23), lacertids like *O.jerdonii* (n = 12) and *O. beddomei* (n = 11), and agamids like *C. versicolor* (n = 15), with single encounters of *M. rouxii* and *N. naja*. Individuals of *E. carinatus* were only encountered on the LP and individuals of *H. albofasciatus* were more abundant on the LP compared to the MO. Species like *E. carinata*, *E. allapallaensis* and *O. beddomei* were only found in the orchards, with *C. versicolor* more abundant in the orchards with a few individuals in the plateaus (Fig. 4). The multinomial species classification method classified the species *H. albofasciatus*, *O. jerdonii* and *Echis carinatus* as specialist, *O.beddomei*, *E. allapallensis*, *C. versicolor* and *E. carinata* as generalists and *H. cf. varadgiri* as ‘too rare’ since there were very few observation of these species

**Table 1:** Location of the sites.

| Sr. no | Sites | Site code | Latitude | Longitude |
| --- | --- | --- | --- | --- |
| 1 | Devache Gothane | Plateau 1 | 16.6307 | 73.4159 |
| 2 | Rundhe | Plateau 2 | 16.7296 | 73.3735 |
| 3 | Katradevi | Plateau 3 | 16.52906 | 73.3916 |
| 4 | Barsu | Orchard 1 | 16.6524 | 73.4438 |
| 5 | Nanar | Orchard 2 | 16.5364 | 73.4355 |
| 6 | Ambelkarwadi | Orchard 3 | 16.6801 | 73.4164 |

The Shannon’s entropy index indicated that species composition of orchards (*H’* = 1.797144) were significantly more heterogeneous compared to the plateaus (*H’* = 1.317345) (Hutcheson t-statistic = 5.2389, df = 215.29, p-value = <0.0001) (Fig. 2). The composition of the reptile communities found on the orchards were different from the reptile communities found on the plateaus. The Whittaker’s beta diversity values were more between sites across the two habitats and less within sites across habitats, indicating species turnover (Table 2). The dendrogram plotted using the Whittaker’s beta diversity dissimilarity matrix formed two major clusters, one with the three orchards and other containing the three plateaus, indicating the different composition of species between habitats (Fig. 3).

**Figure 2:**
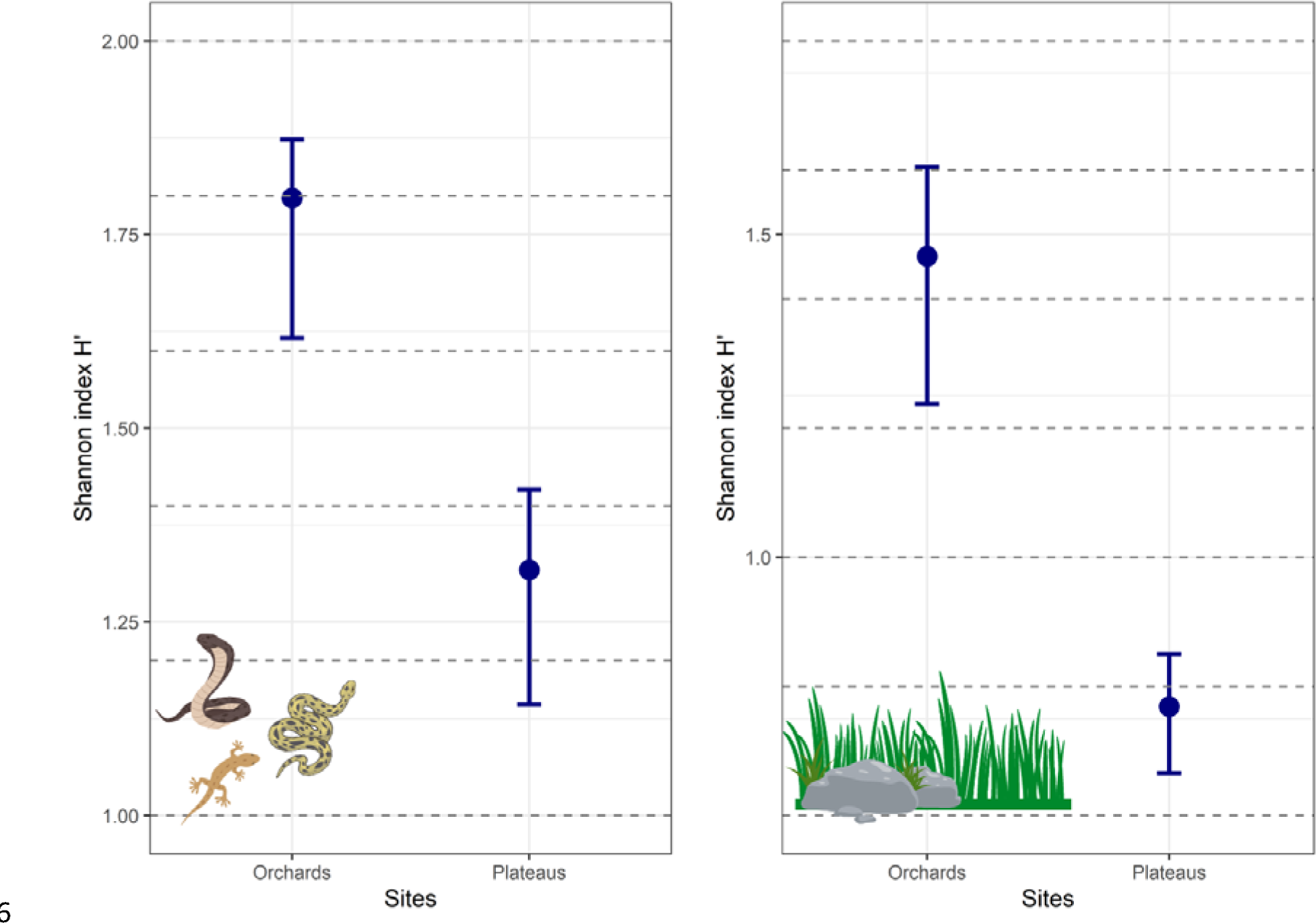
Shannon entropy values for reptile species diversity. Shannon entropy values highlighting the heterogeneity of the microhabitats occupied by reptiles in orchards and plateaus. Error bars indicate 95% confidence interval from bootstrap sampling.

**Figure 3:**
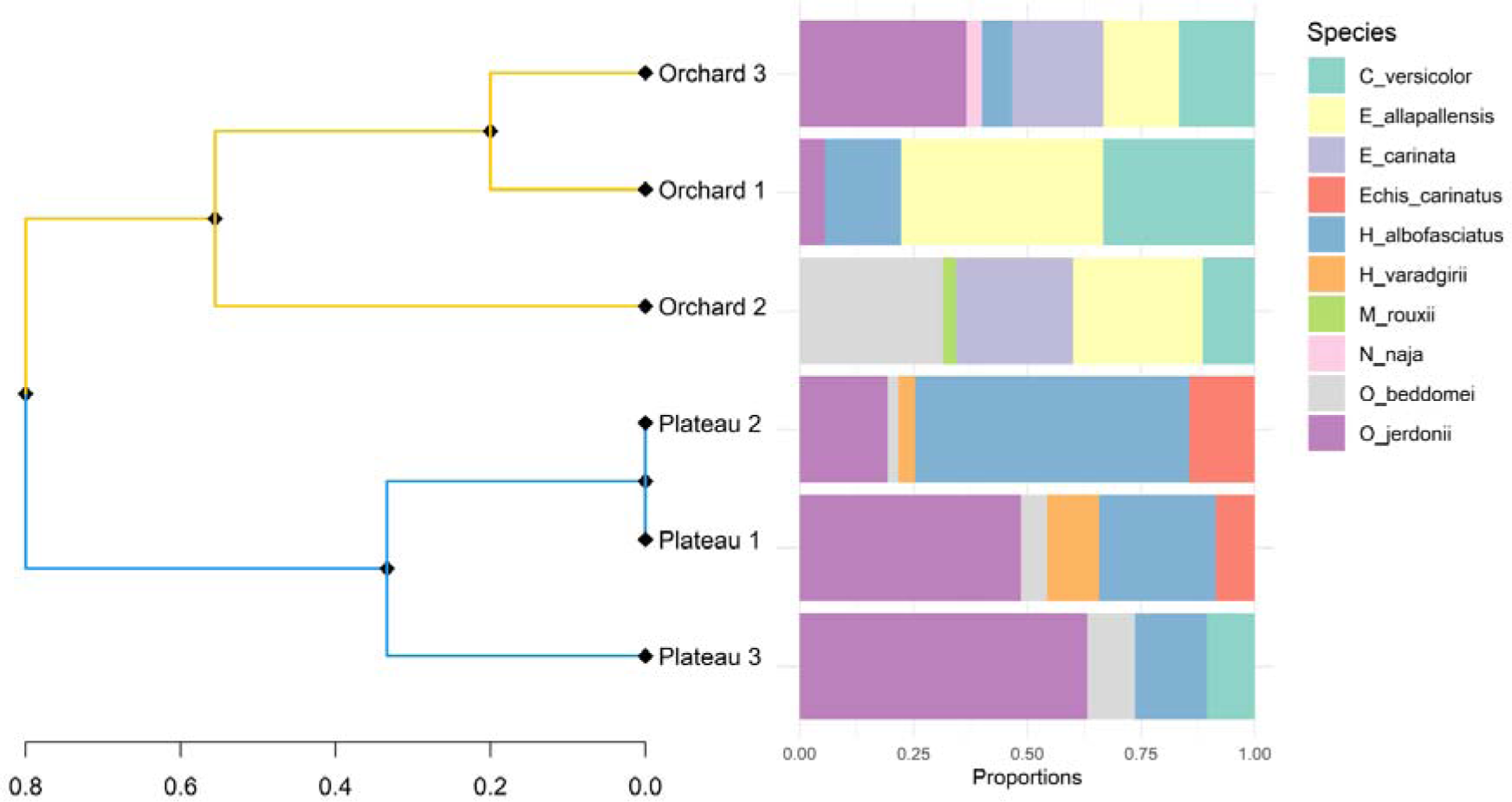
Dendrogram plotted using Whittaker’s beta diversity values, highlighting two major clusters. The stacked-bar plot indicate the proportion of species found in each sites.

**Figure 4:**
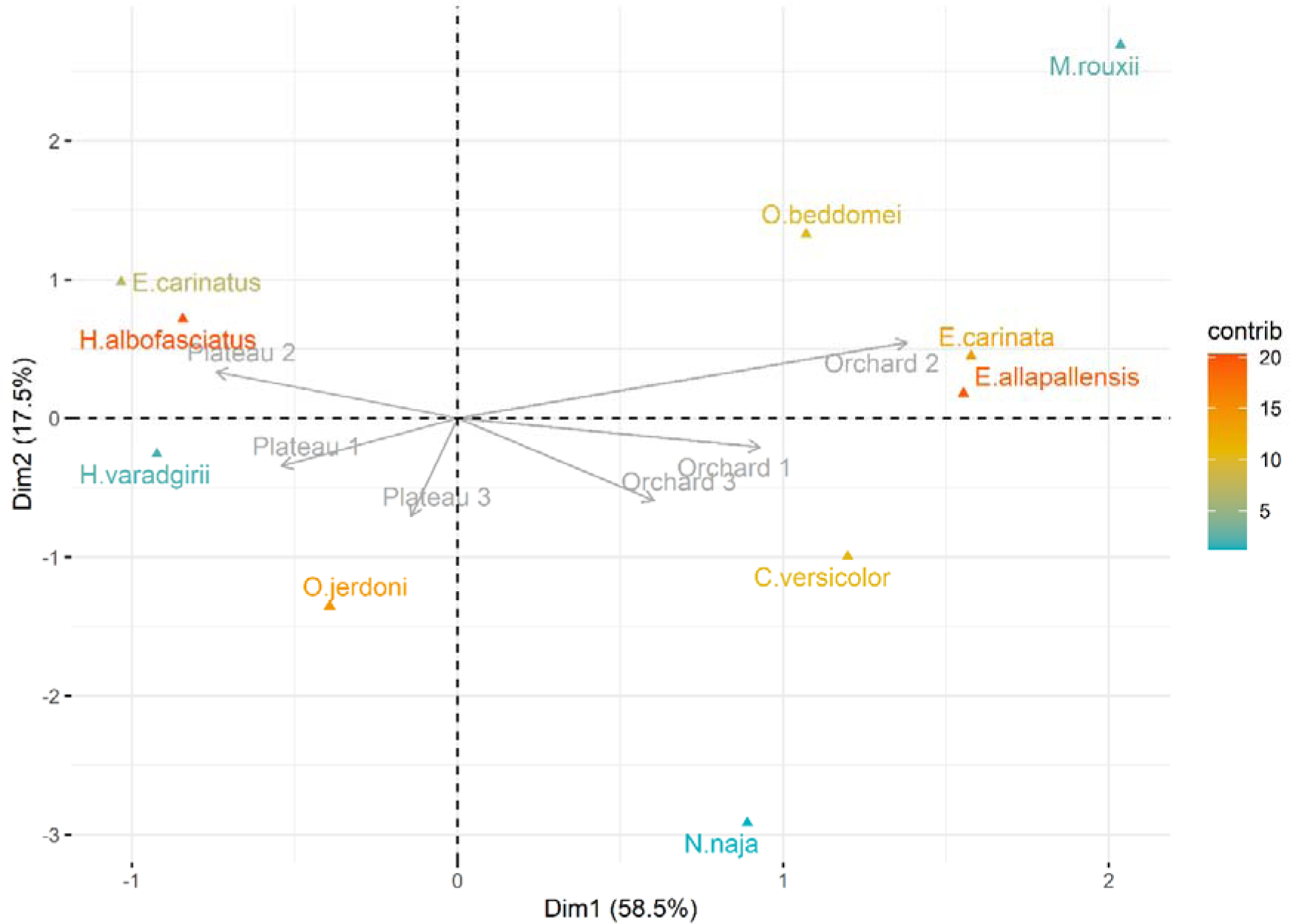
A CA plot showing the distribution of the species within sites

**Figure 5:**
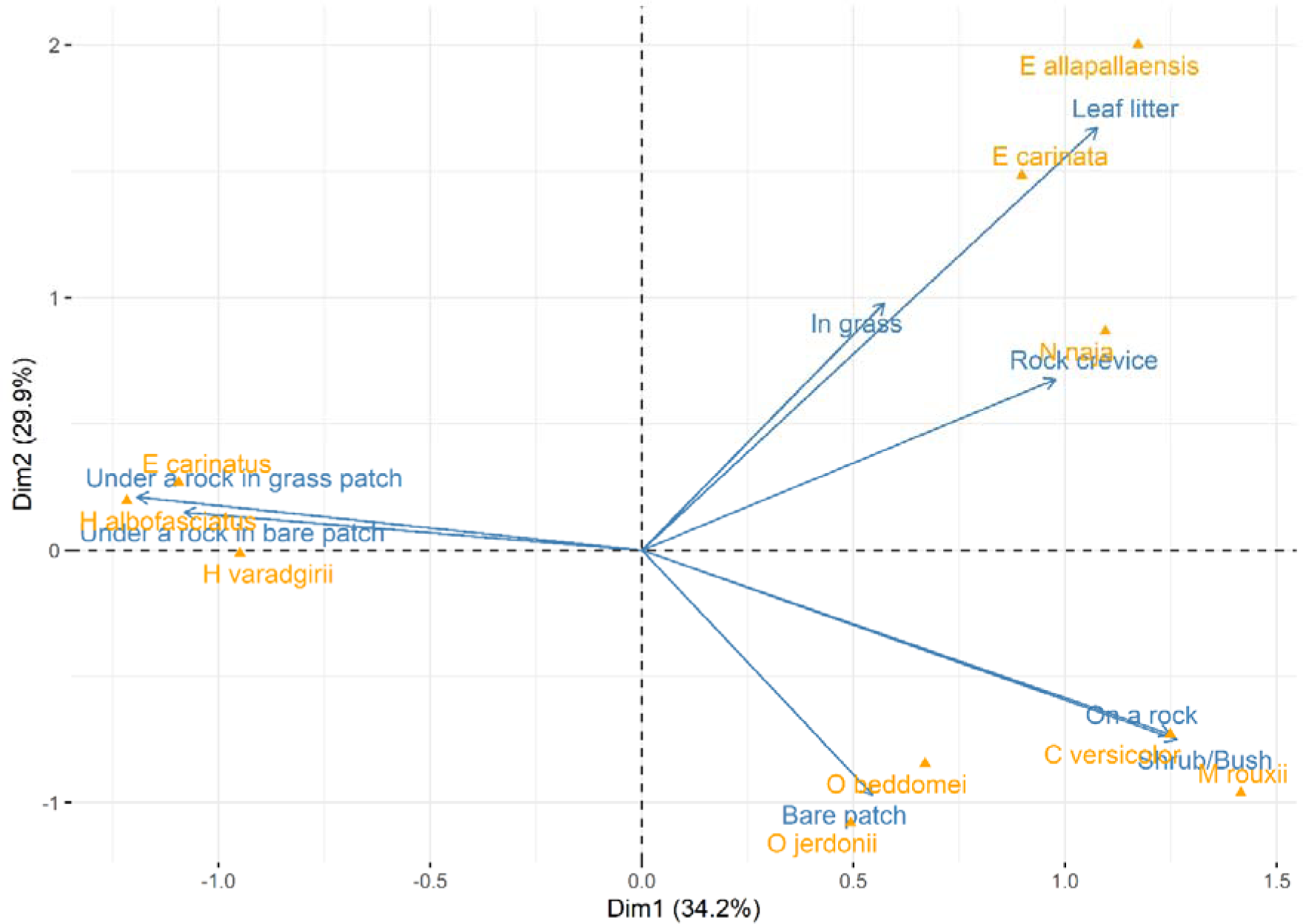
A CA plot showing the microhabitats occupied by reptiles in the orchards and plateaus.

**Table 2:**
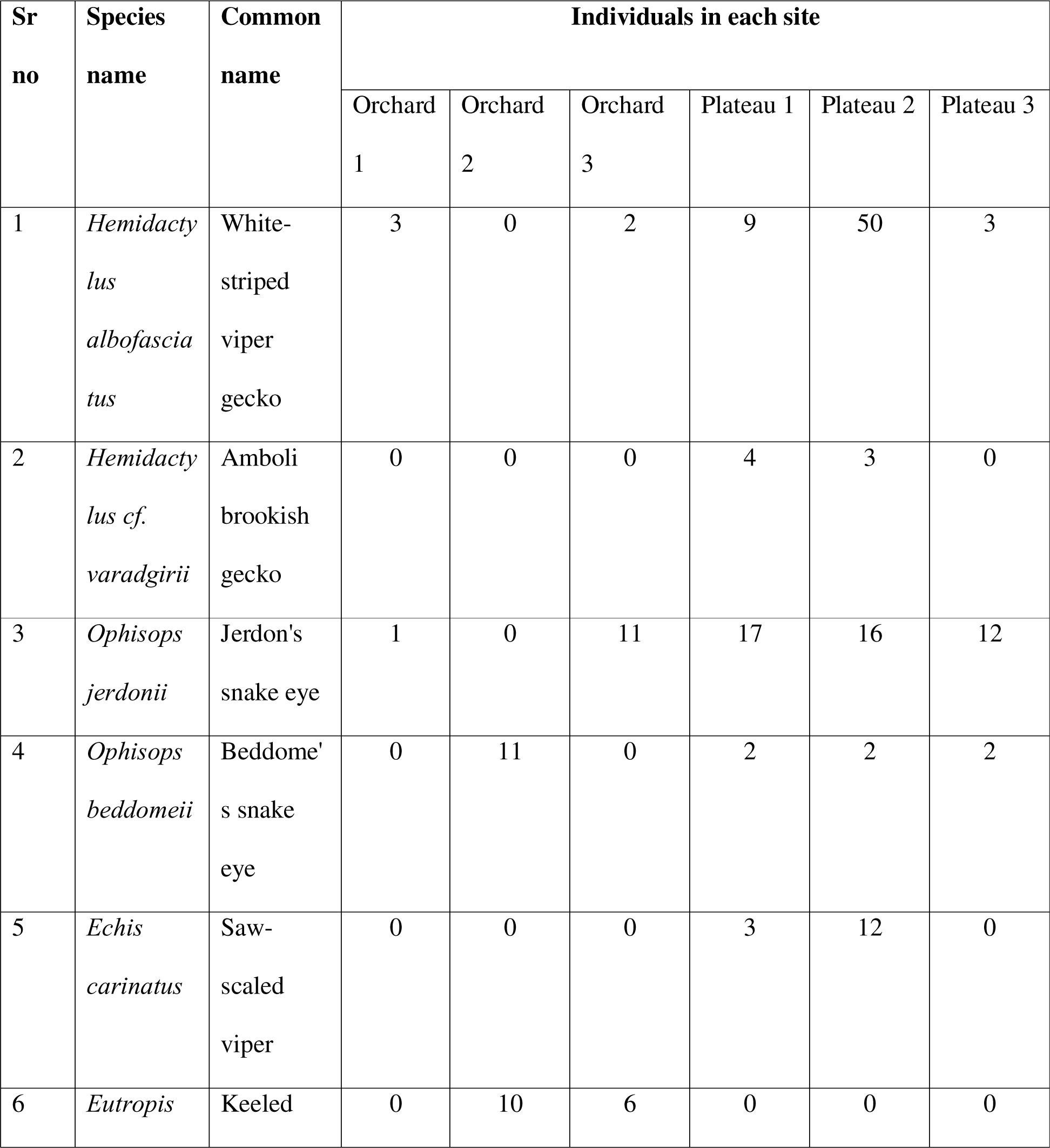
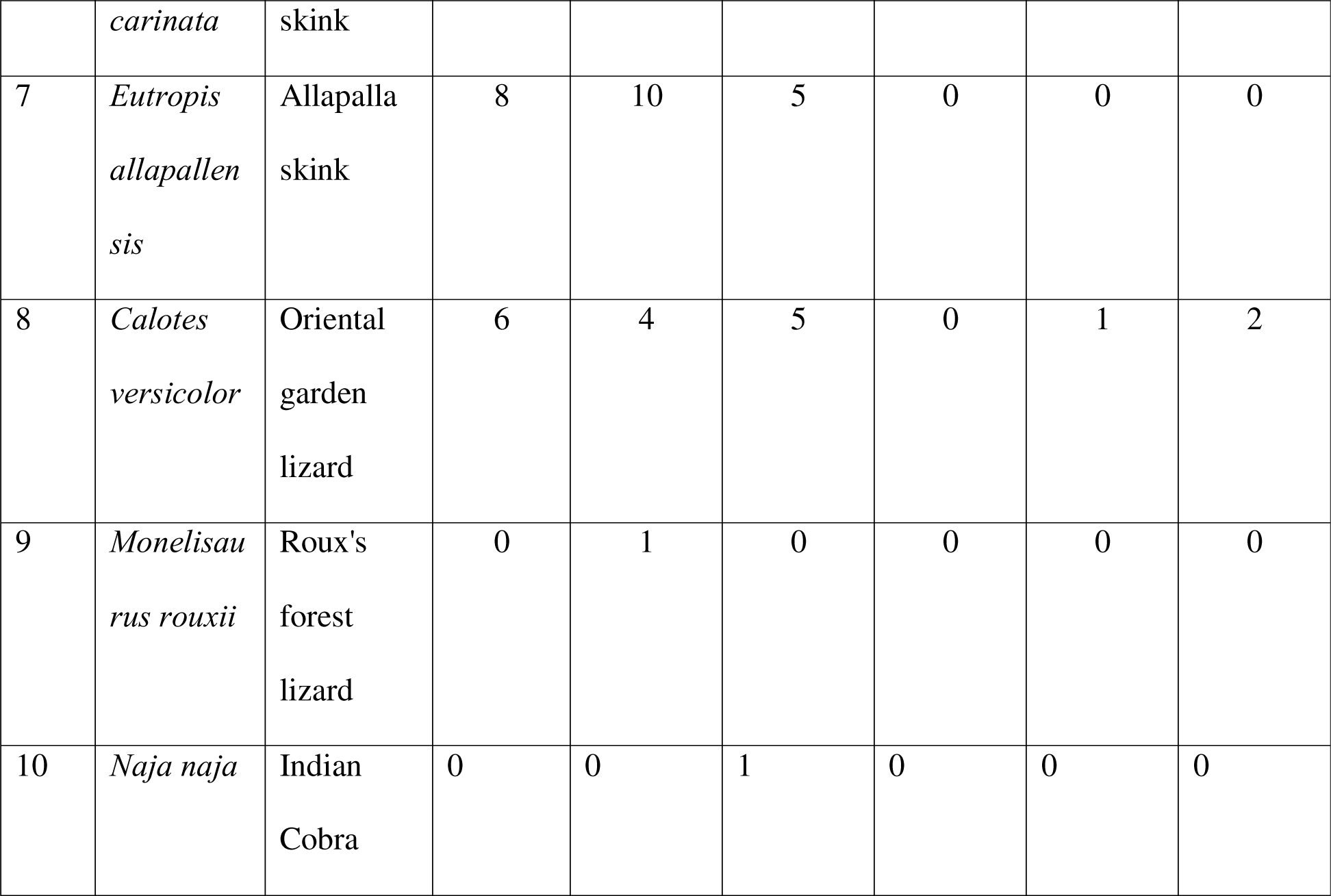
Distribution of species across sites.

**Table 3:**
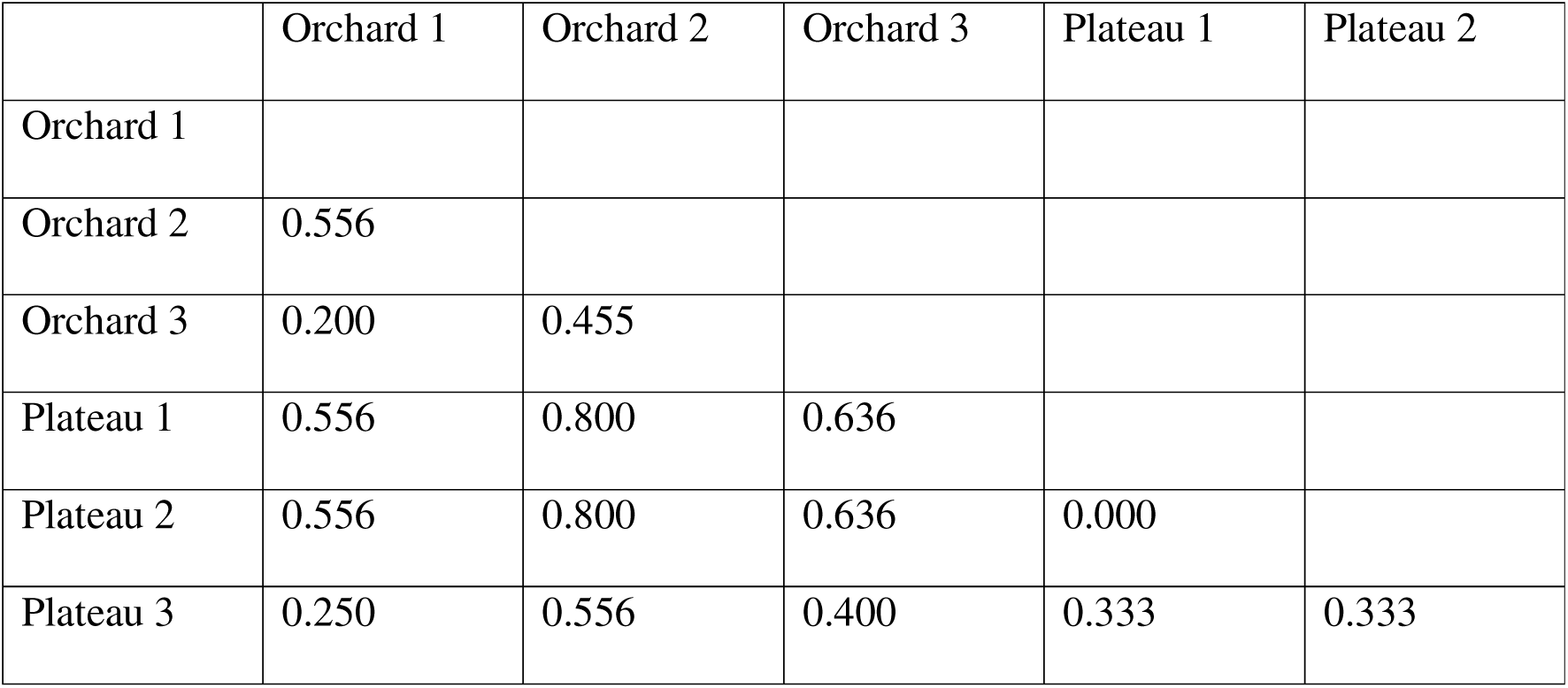
Dissimilarity matrix of beta-diversity values.

|  | Orchard 1 | Orchard 2 | Orchard 3 | Plateau 1 | Plateau 2 |
| --- | --- | --- | --- | --- | --- |
| Orchard 1 |  |  |  |  |  |
| Orchard 2 | 0.556 |  |  |  |  |
| Orchard 3 | 0.200 | 0.455 |  |  |  |
| Plateau 1 | 0.556 | 0.800 | 0.636 |  |  |
| Plateau 2 | 0.556 | 0.800 | 0.636 | 0.000 |  |
| Plateau 3 | 0.250 | 0.556 | 0.400 | 0.333 | 0.333 |

### 3.2. Microhabitats use across habitats

The microhabitats and substratum occupied by reptiles in the orchards (*H’* = 1.4662522) were more significantly heterogeneous compared to that occupied in the plateaus (*H’* = 0.7687774) (Fig. 2) (Hutcheson t-statistic = 6.7687, df = 128.72, p-value < 0.0001). The species found on the plateaus like *H. albofasciatus*, *H. varadgirii* and *E. carinatus* occupied the empty spaces between the loose rocks present on the bare-patches with very few individuals found under rocks in grass with little soil. Other species like *O. jerdonii* were found actively foraging on bare-ground, without any grass-cover. The few individuals (n = 3) of *C. versicolor* encountered on the plateaus were found on a few the scattered Lantana bushes. In the orchards, skinks like *E. carinata* and *E. allapallensis* were found actively foraging in the leaf-litter, while lacertids (*O. jedonii* and *O. beddomei*) were found on bare ground. *C. versicolor* occupied the shrubs, bushes and rocks used to build bunds around individual trees in the orchards, as perches (Fig. 6).

**Figure 6:**
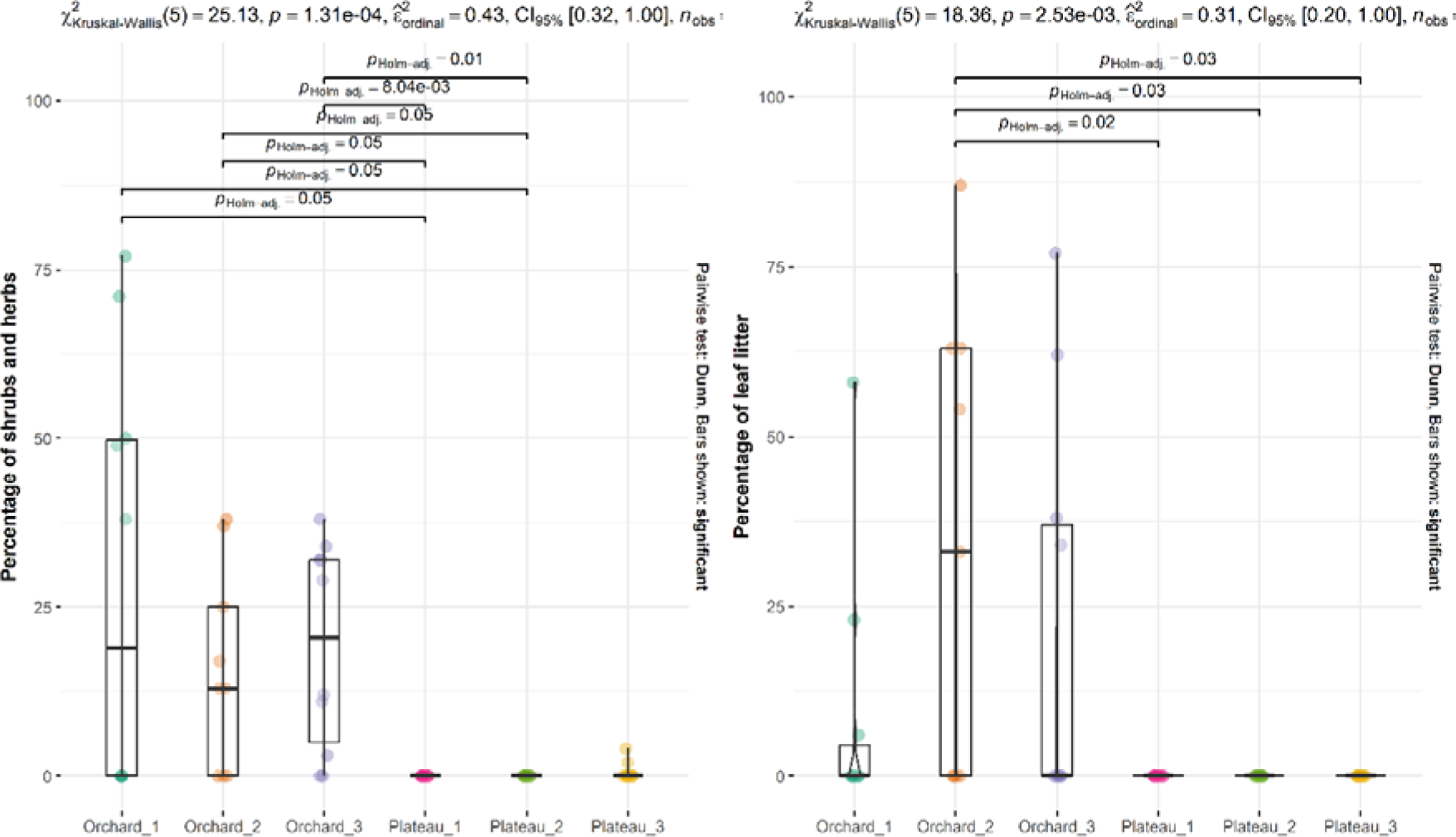
Percentage of shrubs and herbs, and leaf-litter across sites. The bar indicate the adjusted p-values for Dunne’s post-hoc test.

### 3.3. Change in substratum across sites

We found no significant difference in the percentage of bare patch (Kruskal-Wallis chi-squared = 7.4614, df = 5, p-value = 0.1885) and grass cover (Kruskal-Wallis chi-squared = 3.9502, df = 5, p-value = 0.5566) across sites (Fig. 6). However, there was significant difference in the percentage of leaf-litter (Kruskal-Wallis chi-squared = 18.358, df = 5, p-value = 0.00253) and small herbs and shrubs (Kruskal-Wallis chi-squared = 25.133, df = 5, p-value = 0.0001313) across sites (Fig. 7).

**Figure 7:**
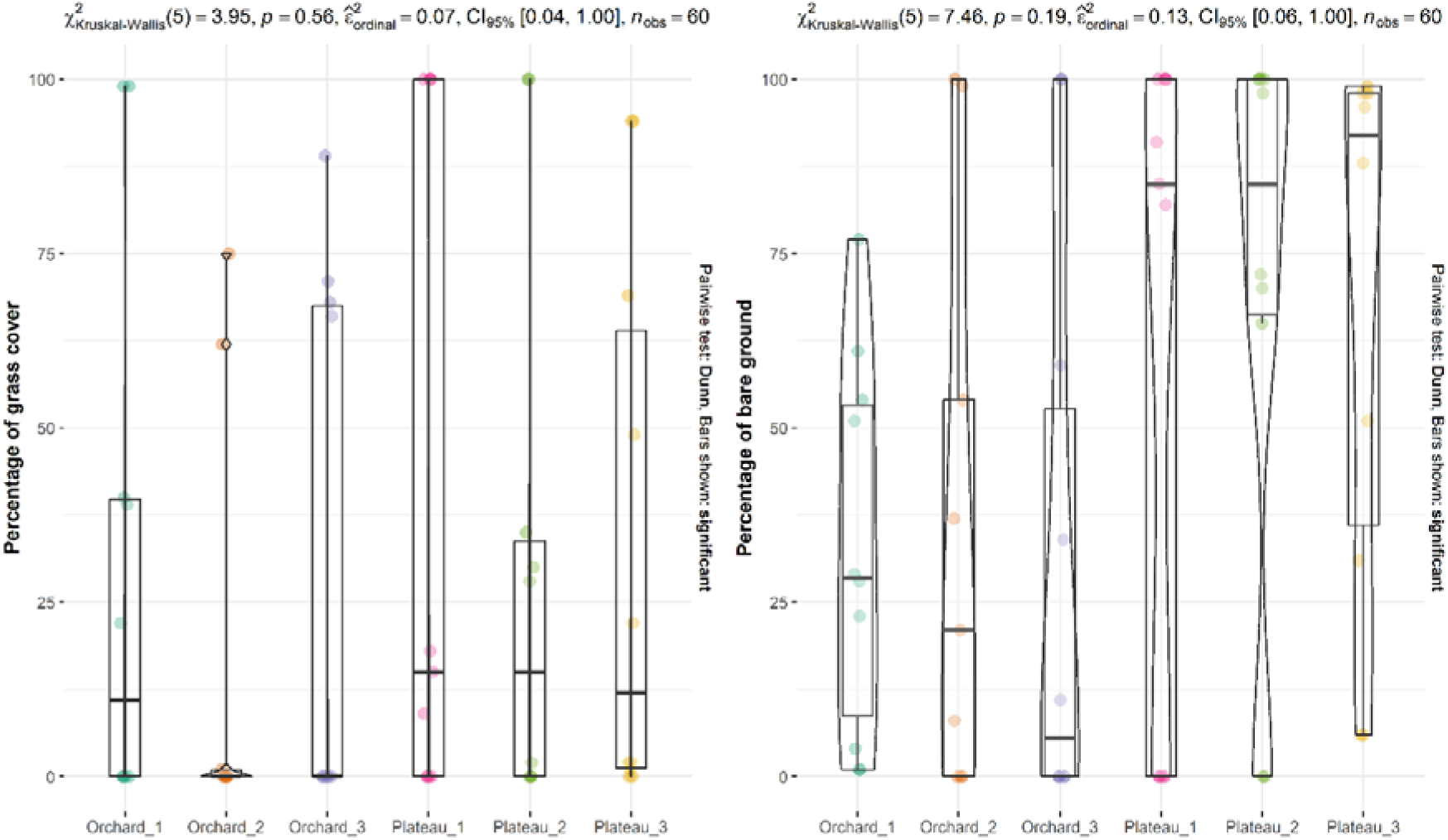
Percentage of substratum (Grass and bare-patches) across sites showing no significant changes.

## 4. DISCUSSION

Considering the rapid change in land-use patterns on the lateritic plateaus in Konkan, it is important to understand the effects of these changes on the bio-diversity for effective management of the landscape and conservation. Ratnagiri and Sindhudurg districts in Maharashtra, India are one of the leading producers in the Alphonso variety of mangoes (Burondkar et al., 2018). The locals here have started converting the plateaus in to mango monocultures. Hence, in this study we examined the effects of converting the plateaus into orchards on the reptile communities. We sampled reptile in both habitat along with their microhabitat preferences and our findings revel important insights into the diversity, distribution and habitat preferences of reptiles in both habitats.

### 4.1. Sampling and Study design

Generally, while evaluating the impact of human activity on biodiversity, two approaches are used, space-for-time (SFT) substitution and before-after control-impact (BACI). In SFT the evaluation is done after the main disturbance event have already taken place and compared to spatial reference sites where the human impact has not yet occurred. On a contrary, in BACI the assessment is done prior to the disturbance and again, after the disturbance event has occurred. The BACI approach has proven to be more efficient compared to the SFT (França et al., 2016). In this study we used the SFT, since BACI are more resource and time consuming. However, irrespective of the approach, we were able to capture the effect of habitat alteration on these reptile communities.

The reptiles found on the LP lateritic plateaus like *H. albofasciatus* and *E. carinatus* only occupy microhabitats like loose spaces under rocks to during the day, making them easy to detect by turning rocks during the days. Hence, we carried out sampling during 1500 hrs – 1800 hrs using active search method. The scope of the study was to examine how the composition of species differed when the plateaus are converted in to orchards and not to catalogue all the reptile species found in the landscape. Hence, although we opportunistically spotted widespread and generalist species like *Ptyas mucosa*, *Amphiesma stolatum*, and *Fowlea piscator* and *Riopa guentheri* outside the belt transects in the orchards, they were not included in the analysis.

### 4.2. Reptile composition and microhabitat preference across habitats

There was a significant difference in heterogeneity in species composition between the plateaus and the orchards. The calculated values for *H’* indicate that the orchards were more heterogeneous in species composition compared to the plateaus. We found species like the *H. albofasciatus*, *E. carinatus* and *O. jerdonii* with a few individuals of *C. versicolor* in the plateaus, similar to the observations made by Gaikwad et al. 2010, and species like *E. carinata*, *E. allapallensis*, *C. versicolor* and *O. beddomei*, with few individuals of *O. jerdonii* and *H. albofasciatus*, and single observations of *N. naja* and *M. rouxii*, in the orchards.

### 4.3. More heterogeneity in the species composition in the orchards, should the plateaus be converted in to orchards?

When forests are altered into plantations reptile species experience a species-turnover rate (VegalAgavo et al., 2021). Species which tend to occupy heterogeneous niches, ‘generalist species’ are less vulnerable to habitat alteration compared to specie which occupy homogenous niches, ‘specialist species’. Habitat specialist and generalist species are distributed across habitats, depending on the structure of the habitat (IglesiaslCarrasco et al., 2023). Although the heterogeneity of the species composition was more in the orchards compared to the plateaus, generalist species were more abundant in the orchards compared to the plateaus. On a contrary, specialist species were more abundant in the plateaus compared to the orchards. For example - *H. albofasciatus*, endemic to the LP in Konkan, Maharashtra and are categorized Vulnerable (VU) under the IUCN read list of endangered species (IUCN, 2011), this species was abundant on the plateaus compared to orchards. Although, *E. carinatus* are widespread in India, we found them only on the plateaus. Hence, converting the plateaus into orchards causes a species turnover, with the orchards harbouring more generalist species compared to specialist species.

### 4.4. Micro-habitat and distribution of reptiles

Similar to the species composition, there was a significant changes in the microhabitat occupied by the reptiles. The microhabitats occupied by the reptiles on the orchards were more heterogeneous compared to the plateaus. Hence, the more heterogeneity in species composition in the orchards compared to plateaus could be explained by this. The availability of loose rocks on the plateaus have been proven to be an important factor to determine the presence of *H. albofasciatus* and *E. carinatus* in the plateaus, orchards and paddy fields (Jithin et al., 2023). We found similar trends in our study with *H. albofasciatus* and *E. carinatus* only occupying these loose rocks on the plateaus as well as the orchards. Addition of trees to the landscape increased the leaf-litter, adding more new niches for reptiles to occupy. Additionally, the bunds made around each Mango tree using rocks sourced from the LP, also provide more open niches for reptiles. Species of skinks like *E. carinata* and *E. allapallensis* are commonly found around human modified landscape and were abundant in the orchards, actively found foraging in the leaf litter and sometimes were detected between these rock crevices. Similar*, C. versicolor* is a widespread in India and found in human-modified landscapes like plantations and human settlements. We found this species more abundant in the orchards compared the plateaus. In the orchards, *C. versicolor* occupied the branches of Mango trees and the bunds made around trees with rocks as perching sites. In the plateaus, they occupied the invasive bushes of *Lantana camara* for perching and basking, and were never found on the ground. The orchards offered more microhabitats for the reptiles to occupy compared to the plateaus.

Global trends in conversion of natural habitats to monocultures and plantations have proven that, loss of biodiversity is not linked to loss of particular groups with specific body size, but the instead maintained similar size disparity compared to forests (IglesiaslCarrasco et al., 2023). We found contrasting results in our study. Species like *H. albofasciatus* has adapted certain traits like small-body size (Largest – Snout-vent length (SVL) - 35mm), which helps them to occupy these spaces under loose rocks to protect themselves from predators and extreme weather conditions during the day. Similarly, *E. carinatus*, with small body size also occupied similar microhabitats as *H. albofasciatus* and were also detected under the same rock. On the contrary, species in like *E. carinatus*, *E. allapallensis*, *C. versicolor*, *N. Naja* and *M. rouxii*, found in the orchards had relatively larger body sizes (Larger SVL). This could be due to the additional niches provided by the Mango trees in the orchards, favouring species with larger body sizes. Similar, there could be other morphological traits which might be restricting the generalist species to the orchards. Further studies could be carried out, examining the functional diversity of the reptile communities in the landscape in order to address this research gap.

### 4.5. Conservation of the lateritic plateaus

The plateaus face threats such as change in land-use patterns (converting the plateaus into mango orchards, cashew plantations, rock quarries and agriculture), and other anthropogenic threats such as deliberate fires and removal of rocks which act as a crucial microhabitats for small invertebrates and vertebrates found here (Watve, 2013). Additionally, in the 1800’s *E. carinatus* was considered as a health hazard. The British government, in India ran campaigns for culling individuals of *E. carinatus*. A total of 1,354,330 individuals were killed and the locals were compensated against each dead individual been returned to the officials. Later in the 1900’s, The Haffkine institute, Mumbai started collecting live specimens of *E. carinatus* from Konkan, sourced from the lateritic plateaus for collecting venom for the production of anti-venom. Around 59,766 individuals were collected from Sindhudurg and Ratnagiri district respectively between 1971 and 1991(Sengupta et al., 1994). A few projects like the Jaitapur Nuclear power plant and the Nanar oil refinery have also been proposed on these plateaus, covering 1000’s of acres of lands.

Although a few plateaus in the Ratnagiri and Sindhudurg districts have been proposed as Biodiversity Heritages Sites, majority of the plateaus are still not legally protected. Hence, it is necessary to come up with a new conservation model which will conserve all the plateaus and simultaneously allow the local communities the rights over their land and conserve biodiversity at the same time.

## ACKNOWLEDGEMENT

We would like to express my heartfelt gratitude to Dr Aparna Watve for her generous financial support, which made our fieldwork possible. We would like to thank Dr Vivek Cyriac for his valuable comments on the manuscript. PA would also like to extend my sincere appreciation to Mrs. Christi Sylvia for her invaluable insights and comments on the study design. PA is deeply thankful to Mr. Kailash Amberkar for generously providing his vehicle for our transit between study sites. PA would like to acknowledge and thank Mr. Anirudh Nippani and Mr. Yash Surve for their dedicated participation in the surveys and their constant support in the field. Their contributions have been invaluable to the success of our research project.

## AUTHOR CONTRIBUTIONS STATEMENTS

PA and RM conceived the idea and designed the methodology. PA collected and analysed the data, and led to the writing of the manuscript. RM critically reviewed the draft and gave final approval for publication.

## SUPPLEMENTARY FIGURES

**Figure S1:**
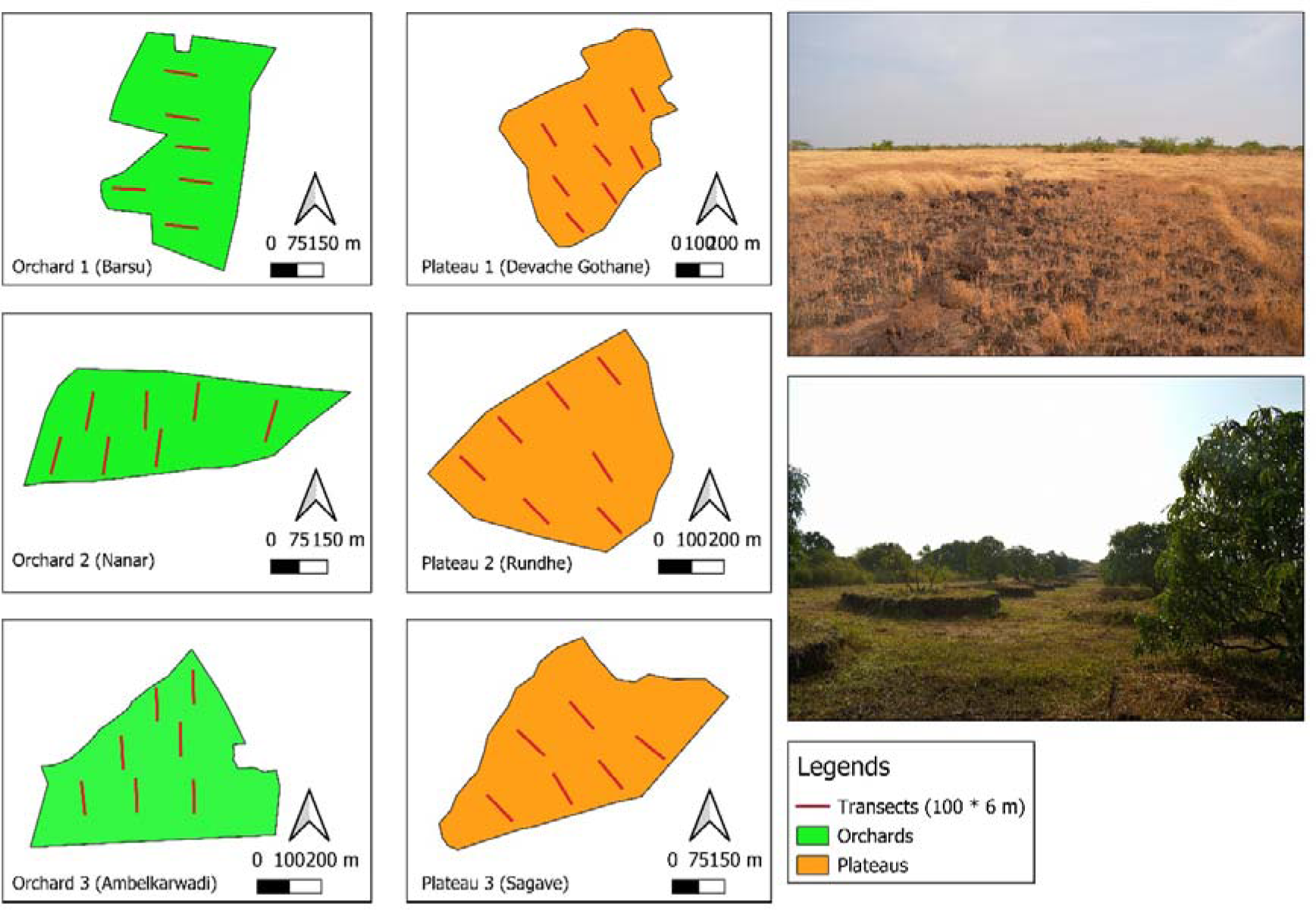
Map of the sites showing distribution of transects, highlighted in red. Right bottom: Photo of an orchard at Ambelkarwadi (Orchard 3). Right top: Photo of a plateau at Rundhe (Plateau 2).

**Figure S2:**
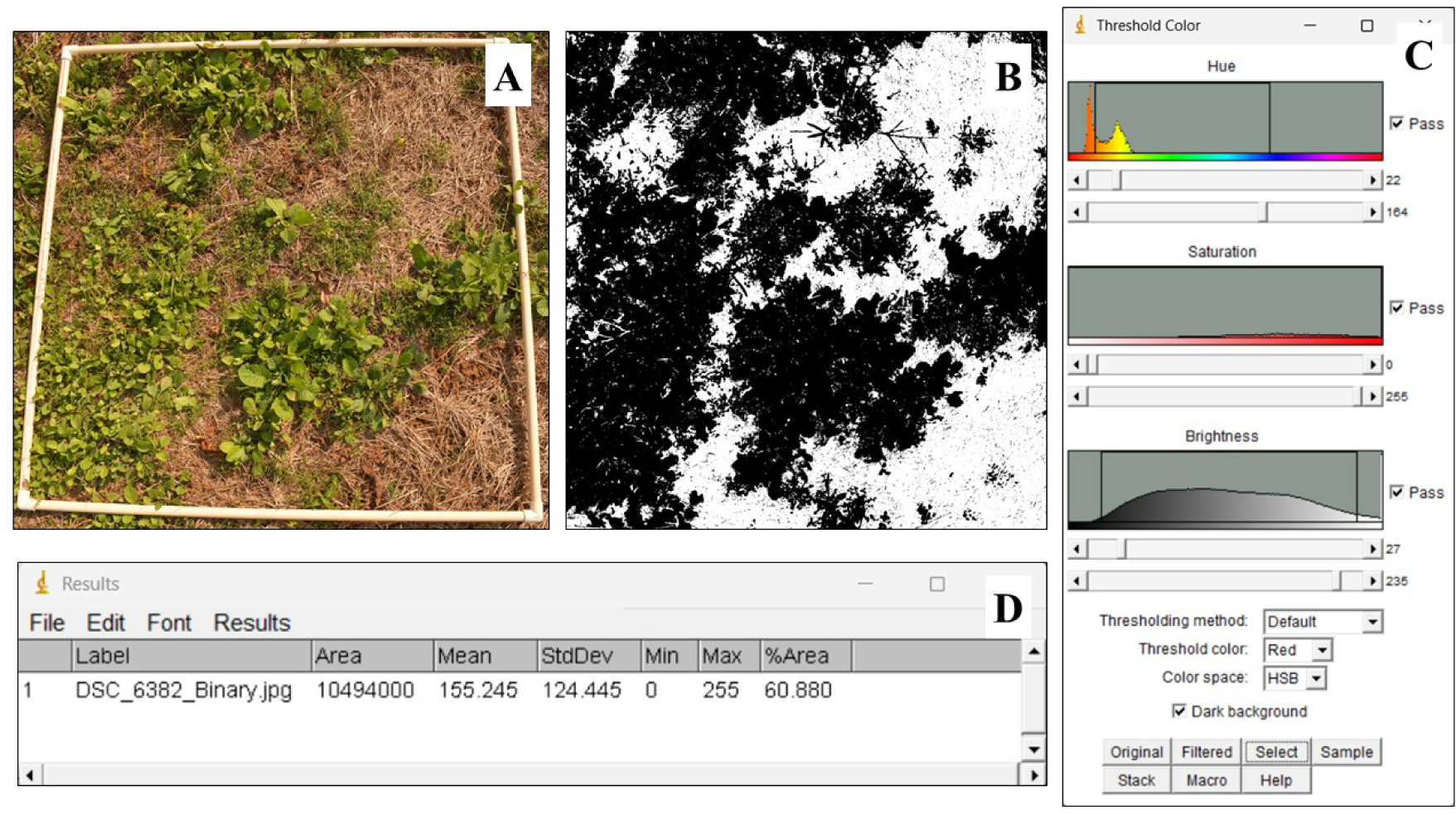
A) Picture of a plot (1×1 meter) from Orchard 3 used for image analysis to estimate the percentage of substratum cover in each site B) Image A converted into a binary image after manually adjusting the colour threshold of each quadrate as shown in figure C and D) the percentage of the green herbs calculated using the ‘Measure’ tool

